# Lamin B1 affects nuclear shape and integrity through chromatin stiffness and not lamin stiffness

**DOI:** 10.64898/2026.08.10.744010

**Authors:** Andy Li, Catherine G. Chu, Nick Lang, Edward J. Banigan, Andrew D. Stephens

**Author notes:** contributing author: Andrew Stephens. co-first authors contributing equally (Andy Li, Catherine G. Chu).

## Abstract

The mechanical properties of the nucleus are critical for maintaining nuclear integrity and function. We previously showed that chromatin dominates short-extension mechanics whereas lamins provide long-extension strain stiffening. To distinguish the roles of lamin isoforms, micromanipulation nucleus force measurements were performed on isolated nuclei from lamin A/C (*Lmna-/-*) and lamin B1 (*Lmnb1-/-*) knockout mouse embryonic fibroblast cells. Lamin A/C knockout does not alter short-extension nuclear stiffness but is essential for strain stiffening at longer extensions. Oppositely, lamin B1 loss reduced short-extension stiffness due to facultative heterochromatin loss while long-extension strain stiffening was slightly increased. Loss of lamin A/C and B1 resulted in similar lamin-chromatin linkers effects as LBR did not change and LAP2β decreased in both. A simulation model of a polymeric lamin shell with stiff lamin A/C and softer lamin B1 subunits can qualitatively recapitulate experimental measurements of lamin knockout cells. Lamin A/C knockout resulted in abnormal nuclear shape but not nuclear blebbing or rupture whereas lamin B1 knockout, similar to other perturbations that cause heterochromatin loss, resulted in increased nuclear blebbing and rupture. This work illuminates the distinct mechanical roles of lamin A/C and B1 in determining nuclear structure and integrity.

## Introduction

Mechanics of the cell nucleus is essential to many core functions of the cell, including maintaining nuclear shape and compartmentalization, migration, and response to external cues via mechanotransduction. Many human afflictions such as aging, heart disease, muscular dystrophy, and many types of cancer present disruption of nuclear mechanical response and protection (Stephens *et al*., 2019a; Kalukula *et al*., 2022). The two main mechanical components of the nucleus are lamins, which form a thin meshwork at the nuclear envelope, and chromatin, which fills the interior. A number of studies have revealed that these two mechanical elements have different roles (Stephens *et al*., 2017; Hobson *et al*., 2020; Bergamaschi *et al*., 2024). Chromatin provides the dominant contribution to mechanical response to small deformations due to its elasticity and the comparative flexibility of shells under axial tension (Banigan *et al*., 2017; Stephens *et al*., 2017, 2018; Attar *et al*., 2025). Due to their short persistence length lamins bend easily (Mahamid *et al*., 2016; Turgay *et al*., 2017), but at longer extensions, the lamina stretches, causing the nucleus to strain stiffen (Stephens *et al*., 2017; Hobson *et al*., 2020). Physical simulation models support this experimental finding (Banigan *et al*., 2017; Hobson and Stephens, 2020). However, the individual contribution of each type of lamin to strain stiffening is not known.

Lamins are intermediate filaments that form a meshwork at the nuclear envelope (Shimi *et al*., 2015). Lamins are split into two classes. A-type lamins, consisting of lamin A and lamin C, are splice variants from the LMNA gene. B-type lamins are read from the LMNB1 and LMNB2 genes. Farnesylation on lamin B causes it to be more proximal to the nuclear envelope while lamin A/C is more internal, since its farnesyl group is cleaved off (Nmezi *et al*., 2019). It is well established that lamin A/C is a major mechanical component of nuclear lamina (Lammerding *et al*., 2004, 2006; Pajerowski *et al*., 2007; Schäpe *et al*., 2009; Swift *et al*., 2013; Stephens *et al*., 2017; Sapra *et al*., 2020; Vortmeyer-Krause *et al*., 2020). Oppositely, lamin B1 has been reported to provide either no stiffness to the nucleus (Lammerding *et al*., 2006; Stephens *et al*., 2017) or to weaken the nucleus (Shin *et al*., 2013), leading to the conclusion that nuclear stiffness due to lamins is proportional to the ratio of lamin A to lamin B (Swift *et al*., 2013; Harada *et al*., 2014). The basis for the distinct mechanical roles of lamin A/C and B1 remain to be determined.

The other major nuclear mechanical component is chromatin and its histone modification state (Schreiner *et al*., 2015; Le *et al*., 2016; Shimamoto *et al*., 2017; Stephens *et al*., 2017; Hobson *et al*., 2020; Nava *et al*., 2020; Bergamaschi *et al*., 2024; Williams *et al*., 2024). Importantly, chromatin-based nuclear rigidity is independent of the contribution of lamin A/C (Stephens *et al*., 2017, 2018). However, lamin B1 depletion results in loss of the facultative heterochromatin marker H3K27me3 (Camps *et al*., 2014; Stephens *et al*., 2018; Vahabikashi *et al*., 2022; Pho *et al*., 2023), suggesting that lamin B1 might indirectly impact chromatin-based nuclear rigidity. Oppositely, loss of lamin A/C has not been shown to impact histone modification state.

Another hypothesized nuclear mechanical component is lamin-chromatin linker proteins (Attar *et al*., 2025). Lamin B1 interacts with Lamin B receptor (LBR, (Olins *et al*., 2010; Odell *et al*., 2026)), which tethers chromatin to the periphery (Alabi *et al*., 2025), and lamin-associated protein 2 β (LAP2β,(Alabi *et al*., 2025; Lewis *et al*., 2026)). Thus, lamin-chromatin linker proteins could underlie the differences between lamin B1 and lamin A/C contributions to nuclear mechanics and shape.

The differential mechanical roles of lamins could provide information on the effect of nuclear stiffness on nuclear shape and integrity. Nuclear blebs are an interphase-based herniation of the nucleus that has decreased DNA density (Bunner *et al*., 2024; Chu *et al*., 2025; Pujadas Liwag *et al*., 2025) and result in nuclear rupture and dysfunction (Vargas *et al*., 2012; Stephens *et al*., 2019b; Stephens, 2020; Kalukula *et al*., 2022). Perturbations that are known to alter mechanics, such as lamin A/C mutations and heterochromatin loss often lead to nuclear blebbing and rupture (Goldman *et al*., 2004; Shumaker *et al*., 2006; Stephens *et al*., 2018, 2019a; Eskndir *et al*., 2025; Manning *et al*., 2025; Pujadas Liwag *et al*., 2025). Lamin B1 loss is one of the first and most studied perturbations that increases nuclear blebbing and rupture (Lammerding *et al*., 2006; Vargas *et al*., 2012; Hatch and Hetzer, 2016; Chen *et al*., 2018; Stephens *et al*., 2018; Pho *et al*., 2023; Alabi *et al*., 2025). However, the mechanical basis of lamin B1 loss effects remains to be determined.

To determine the differential roles of lamins in nuclear stiffness, we measured the nuclear spring constant for the short and long extension regimes in lamin knockout cell lines using dual micropipette micromanipulation force-extension measurements. We used immunofluorescence to verify lamin loss and assay facultative heterochromatin levels. We treated wild type cells with the EZH2 methyltransferase inhibitor Tazemetostat to establish the mechanical role of facultative heterochromatin. We also measured lamin-chromatin linking proteins LBR and LAP2β to determine their possible roles. To better understand our findings, we compared our experimental results with molecular dynamics simulations of a shell-polymer model of the cell nucleus. This model demonstrates how the loss of B-type lamins may increase the elasticity of the nuclear lamina. Finally, we show the resulting functional consequences of chromatin-and lamin-based nuclear stiffness on nuclear shape and compartmentalization.

## Results

### Knockout of lamin A/C and B1 have differential effects on the lamina and heterochromatin

To determine the differential roles of lamin A/C and B1, we used stable knockout cell lines in MEFs. Loss of lamin A/C (*Lmna-/-*) and lamin B1 (*Lmnb1-/-*) were previously generated through stable knockout cell lines (Kim *et al*., 2011; Guo and Zheng, 2015). We also include a EZH2 histone methyltransferase inhibitor, Tazemetostat (Knutson *et al*., 2014), as a control for a perturbation that does not alter lamin levels but alters facultative heterochromatin levels.

To verify loss of individual lamins in these cell lines, we conducted immunofluorescence. Immunofluorescence of lamin A/C antibody labeling in *Lmna-/-* measured a near complete loss of lamin A/C (**Figure 1A**). *Lmna-/-* cells also showed a twofold increase in lamin B1 levels. Loss of lamin B1 in *Lmnb1-/-* was confirmed via immunofluorescence of lamin B1 antibody labeling, but without a change in lamin A/C levels (**Figure 1A**). Treatment of wild type cells with Tazemetostat did not alter lamin A/C or B1 levels. Overall, we verify loss of either lamin A/C or lamin B1.

**Figure 1.**
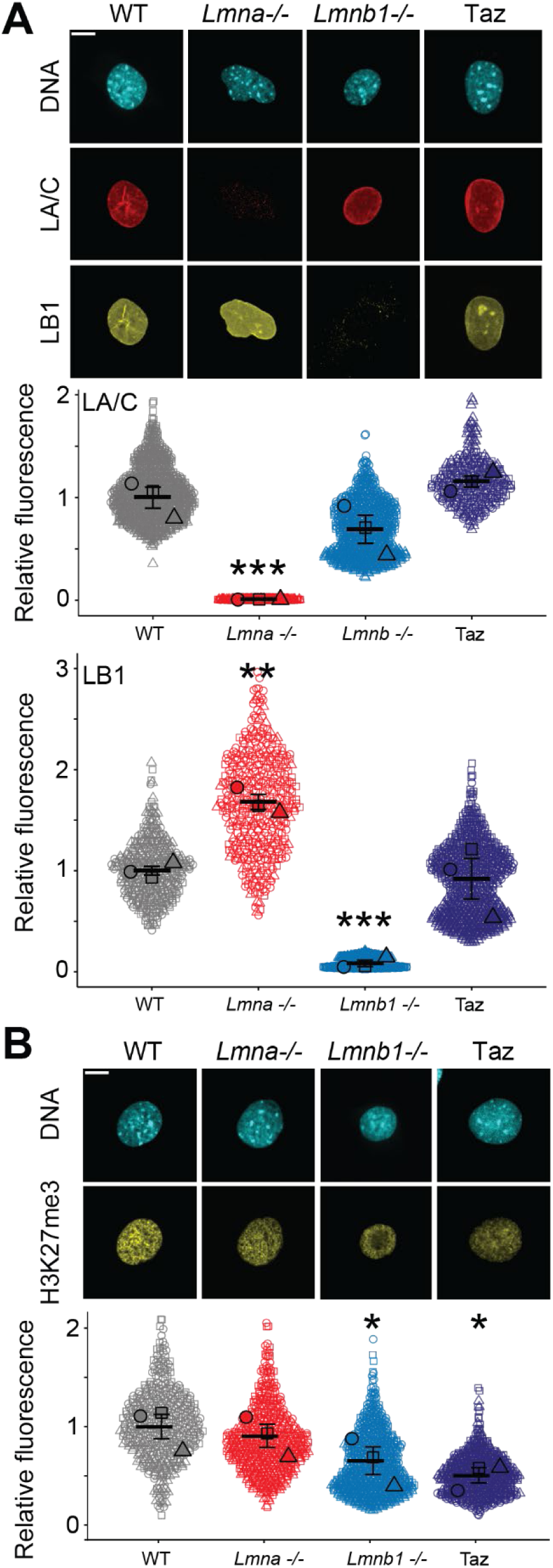
Immunofluorescence measurements of lamin and heterochromatin levels upon lamin A/C knockout, lamin B1 knockout, or Tazemetostat treatment. (A) Example images and superplot graphs of lamin A/C and lamin B1 immunofluorescence in wild type (WT, gray), lamin A/C knockout (*Lmna-/-*, red), lamin B1 knockout (*Lmnb1-/-*, blue), and loss of facultative heterochromatin (Taz, purple). (B) Example images and superplot graphs of facultative heterochromatin marker H3K27me3 intensity in wild type (WT, gray), lamin A/C knockout (*Lmna-/-*, red), lamin B1 knockout (*Lmnb1-/-*, blue), and loss of facultative heterochromatin (Taz, purple). Biological triplicates each with n > 100 nuclei. For all panels mean and standard error, p values reported as *P<0.05; **P<0.01; ***P<0.001, or ns denotes no significance. Statistical tests are one-way ANOVA with a post-hoc Tukey test. Scale bar = 10 µm.

Next, we measured for previously reported changes in facultative heterochromatin in these lamin knockout cell lines. Immunofluorescence antibody labeling for facultative heterochromatin marker H3K27me3 showed similar levels in wild type and *Lmna-/-* nuclei (**Figure 1B**), in agreement with other publications (Lewis *et al*., 2026). In *Lmnb1-/-* nuclei, levels of H3K27me3 decreased relative to wild type (**Figure 1B**), recapitulating previous publications (Camps *et al*., 2014; Stephens *et al*., 2018; Vahabikashi *et al*., 2022; Pho *et al*., 2023). Wild type cells were treated with an EZH2 inhibitor Tazemetostat to provide a positive control for decrease in H3K27me3 levels relative to wild type. Tazemetostat treatment decreased H3K27me3 to similar levels as *Lmnb1-/-* cells (**Figure 1B**). For all conditions, levels of the constitutive heterochromatin marker H3K9me3 remained similar to wild type (**Supplemental Figure 1**). Interestingly, MEF *Lmnb2-/-* nuclei did not show a loss of H3K27me3 but instead showed a significant increase (**Supplemental Figure 2**). Therefore, lamin B1 knockout and treatment with Tazemetostat cause a significant decrease in facultative heterochromatin whereas lamin A/C knockout shows no change.

### Lamin A/C is essential for nuclear strain stiffening at long extensions

Dual micropipette nuclear isolation and micromanipulation force-extension measurements provide the unique capability to measure the nuclear spring constant (nN/µm) over different regimes of nuclear extension (Stephens *et al*., 2017; Currey *et al*., 2022). Force-extension for nuclei isolated from living cells is measured by extension of the nucleus with the pull micropipette and the resulting deflection of the other force micropipette, multiplied by the premeasured micropipette bending constant (**Figure 2A inset**). The short-extension regime (< 3 µm) measures the chromatin-based regime while the long-extension regime (> 3 µm) encompasses chromatin plus lamin-based strain stiffening (**Figure 2, A-D**). Thus, the long minus the short regime provides a measurement of lamin-based nuclear strain stiffening (**Figure 2E**). Finally, we can calculate the ratio for the different regimes by dividing the long regime by the short regime nuclear spring constant (**Figure 2F**). Wild type dual micropipette micromanipulation force-extension measured a short-extension chromatin-based nuclear spring constant of 0.55 ± 0.02 nN/µm and long-extension chromatin plus lamin-based spring constant of 1.15 ± 0.03 nN/µm, in agreement with previous publications (Stephens *et al*., 2017; Currey *et al*., 2022; Bunner *et al*., 2026).

**Figure 2.**
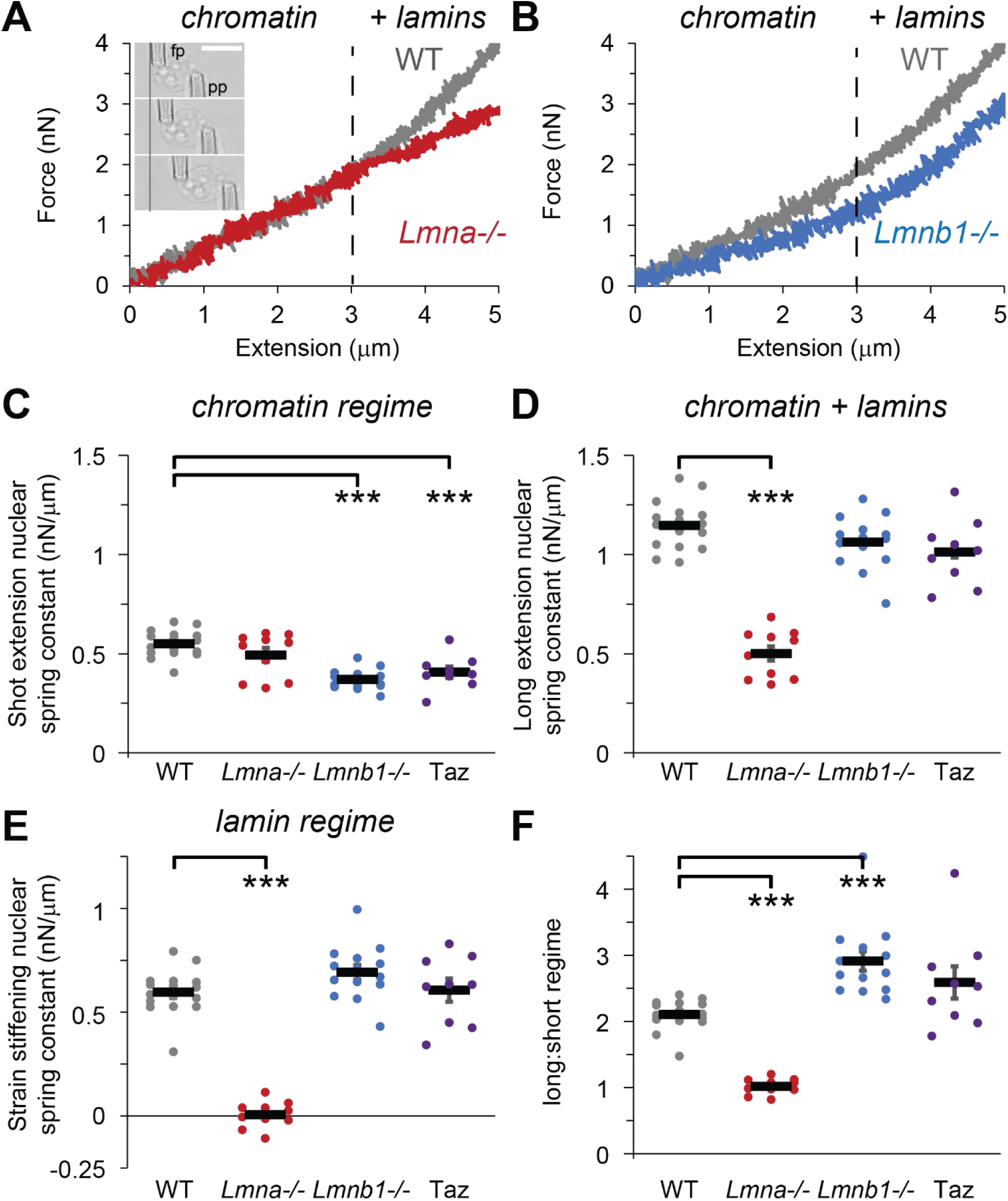
Micromanipulation force measurements reveal differential roles of lamin A/C and lamin B1. (A) Example of micromanipulation isolated cell nucleus force vs. extension measurements for wild type (gray) relative to *Lmna-/-* (A, red) and *Lmnb1-/-* (B, blue). Inset in panel A shows an example of an isolated nucleus suspended between the pull pipette(pp) which when moved causes extension (µm) of the nucleus while deflection of the force pipette (fp) multiplied by its premeasured bending constant provides a measure of force (nN). Short extension (< 3 µm) denotes the chromatin-dominated short extension regime whereas long extension (>3 µm) denotes the addition of lamins that cause strain stiffening. Graph of the nuclear spring constant measured as the slope of the force vs. extension line for the (C) short extension regime measuring chromatin rigidity and (D) long extension regime measuring chromatin and lamin rigidity for wild type (WT, gray), lamin A/C knockout (*Lmna-/-*, red), lamin B1 knockout (*Lmnb1-/-*, blue), and loss of facultative heterochromatin (Taz, purple). (E) Lamin-based strain stiffening nuclear spring constant is derived from subtracting the short extension from the long extension nuclear spring constant. (F) Graph of the ratio of the long/short nuclear spring constant. Biological replicates are individual nuclei n = 16 WT, 10 *Lmna-/-*, 14 *Lmnb1-/-*, 9 Taz. Each nucleus is averaged over 3-4 force-extension measurements meaning totals are WT n > 48, *Lmna-/-* n > 30, *Lmnb1-/-* n > 42, Taz n > 27. For all panels mean and standard error, p values reported as *P<0.05; **P<0.01; ***P<0.001, or ns denotes no significance. Statistical tests are one-way ANOVA with a post-hoc Tukey test. Scale bar = 10 µm.

We previously published that knockdown of lamin A/C in other cell lines does not impact the short-extension regime but is essential for lamin-based strain stiffening at long extensions (Stephens *et al*., 2017; Hobson *et al*., 2020). We aimed to recapitulate this finding in a stable *Lmna-/-* MEF cell line. Micromanipulation force measurements revealed that *Lmna-/-* nuclei were similar to wild type controls in their chromatin-based short-extension nuclear spring constant (**Figure 2, A and C**). This force measurement data agrees with immunofluorescence data showing no change in heterochromatin levels (**Figure 1B and Supplemental Figure 1**). Loss of lamin A/C in MEF *Lmna-/-* cells completely abolished the long-extension strain stiffening regime of the nuclear spring constant. We measured strain stiffening (long – short extension regime) in wild type as 0.60 ± 0.03 nN/µm, which decreased in *Lmna-/-* nuclei to 0.01 ± 0.2 nN/µm (**Figure 2E**). Overall, our new data are in strong agreement with our previously published micromanipulation nuclear spring constants from HeLa and SKOV3 cell lines (Stephens *et al*., 2017; Hobson *et al*., 2020). Thus, lamin A/C is responsible for long-extension nuclear strain stiffening but has no role in the short-extension chromatin-based regime.

### Lamin B1 contributes to nuclear spring constant through facultative heterochromatin maintenance and weakening of lamin strain stiffening

To measure the nuclear spring constant in nuclei lacking lamin B1, we used a MEF *Lmnb1-/-* stable knockout cell line and conducted isolated nucleus dual micropipette micromanipulation force-extension measurements. *Lmnb1-/-* nuclei exhibited a significant decrease in the short extension chromatin-based nuclear spring constant from 0.55 ± 0.02 nN/µm in wild type to 0.37 ± 0.01 nN/µm in *Lmnb1-/-* (**Figure 2, B and C**). This agrees with the loss of facultative heterochromatin levels measured through immunofluorescence (**Figure 1B**). Interestingly, *Lmnb1-/-* nuclei showed no change in the long-extension regime (**Figure 2D**). Furthermore, *Lmnb1-/-* lamin-based strain stiffening response did increase slightly, but not significantly, from 0.60 ± 0.03 nN/µm in wild type to 0.69 ± 0.04 nN/µm in *Lmnb1-/-* (**Figure 2E**). This slight increase in the lamin regime relative to the weaker chromatin regime resulted in significant increase in the long/short regime ratio, from 2.1 ± 0.06 in wild type to 2.9 ± 0.14 in *Lmnb1-/-* (**Figure 2F**). MEF *Lmnb2-/-* nuclei showed no change in short, long, or strain stiffening nuclear spring constant relative to wild type (**Supplemental Figure 2**). Thus, while lamin B1 is part of the nuclear lamina, it has a role in heterochromatin maintenance that impacts chromatin-based nuclear stiffness and has a mild weakening effect on lamin-based strain stiffening.

To determine if loss of facultative heterochromatin alone recapitulates *Lmnb1-/-* nuclear stiffness, we measured the force response of nuclei treated with Tazemetostat, an EZH2 histone methyltransferase inhibitor. Micromanipulation force measurements revealed that Tazemetostat-induced facultative heterochromatin loss resulted in decreased chromatin-based short-extension nuclear spring constant, but no change in lamin-based strain stiffening at longer extensions (**Figure 2C-E**). This data agrees with past measurements of a different histone methyltransferase inhibitor, DZNep, which decreased facultative heterochromatin and the chromatin-based short extension nuclear spring constant, but caused no change in lamin strain stiffening (Stephens *et al*., 2018; Manning *et al*., 2025). Furthermore, the long/short regime ratio in Tazemetostat-treated cells also remains unchanged from wild type, whereas lamin B1 knockout increases significantly (**Figure 2F**). Thus, the Tazemetostat-induced decrease in facultative heterochromatin recapitulates *Lmnb1-/-* decreased nuclear spring constant in the chromatin-based short-extension regime, but not the mild stiffening in the lamin-based strain-stiffening regime.

### Nuclear mechanical differences between loss of lamin A/C and B1 are not due to changes in chromatin-lamin linkers LBR and LAP2β

Lamin A/C and lamin B1 are reported to anchor chromatin through chromatin-lamin linking proteins which could contribute to their mechanical roles. Two important lamin-chromatin tethers are lamin B receptor (LBR) and lamin-associated protein 2 beta (LAP2β) (Solovei *et al*., 2013; Lewis *et al*., 2026). To determine if the nuclear mechanical differences between loss of lamin A/C and lamin B1 are due to changes in chromatin-lamin linkers levels, we measured immunofluorescence within 0.5 µm of the nuclear periphery for LBR and LAP2β (**Figure 3A**). Our measurements revealed that LBR levels remained similar to wild type in both lamin knockout cell lines (**Figure 3B**). This data agrees with recently published work (Lewis *et al*., 2026). Lack of change in LBR was further confirmed by maintenance of peripheral H3K9me3 in both lamin knockout cell lines, which is maintained by LBR (**Supplemental Figure 1**). LAP2β was significantly decreased in both *Lmna-/-* and *Lmnb1-/-* cells relative to wild type (**Figure 3C**). Cells treated with Tazemetostat showed similar outcomes. Overall, loss of lamin A/C or lamin B1 results in similar effects on LBR and LAP2β, suggesting they do not contribute to the mechanism underlying the differences in nuclear strength.

**Figure 3.**
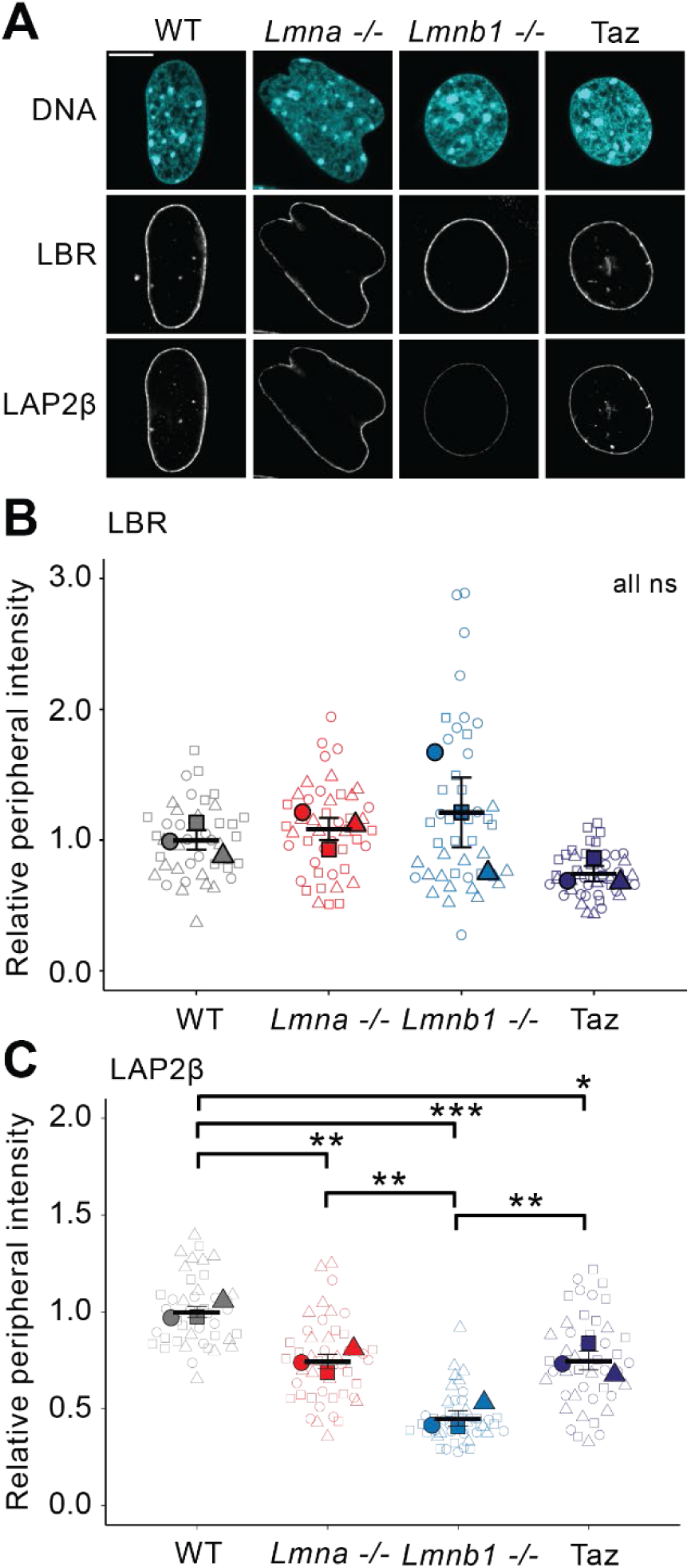
Peripheral intensity of major chromatin-lamin linkers LBR and LAP2β. (A) Example confocal images of DNA via Hoechst along with lamin-chromatin linkers LBR and LAP2β immunofluorescence in wild type (WT), lamin A/C knockout (*Lmna-/-*, red), lamin B1 knockout (*Lmnb1-/-*, blue), and loss of facultative heterochromatin (Taz, purple). Superplot graphs of (B) LBR and (C) LAP2β in wild type (WT), lamin A/C knockout (*Lmna-/-*, red), lamin B1 knockout (*Lmnb1-/-*, blue), and loss of facultative heterochromatin (Taz, purple). Biological triplicates each with n > 15 nuclei. For all panels mean and standard error, p values reported as *P<0.05; **P<0.01; ***P<0.001, or ns denotes no significance. Statistical tests are one-way ANOVA with a post-hoc Tukey test. Scale bar = 10 µm.

### A composite polymer shell model for the lamina suggests different architectures for lamins A/C and B1

We next sought to conceptually understand how depletion of different protein components of the nuclear lamina—lamins A/C and B1—could lead to qualitatively different effects on nuclear stiffness. We particularly focused on explaining the seemingly paradoxical observation that depletion of lamin B1 may stiffen the lamina even though it partially composes this protein meshwork (**Figure 2**; (Shin *et al*., 2013)). To this end, we adapted an established Brownian dynamics simulation model of nuclear mechanical response (See Methods) (Banigan *et al*., 2017; Stephens *et al*., 2017; Strom *et al*., 2021).

In the model, chromatin is treated as a crosslinked polymer inside of a polymeric shell representing the lamina. Chromatin and the lamina are mechanically coupled to elicit mechanical response from both chromatin and the lamina (Attar *et al*., 2025). Monomeric subunits within the polymeric shell were connected by springs, thus forming a meshwork of nodes that resists stretching forces. The model incorporates the hypothesis that lamins A/C and B1 form structures with different microscopic elasticities. Thus, a fraction (25%, 50%, or 75%) of monomeric lamina subunits were randomly selected to have a smaller spring constant. These monomers are referred to as lamin B, while the remaining lamina subunits are referred to as lamin A (**Figure 4A**).

**Figure 4.**
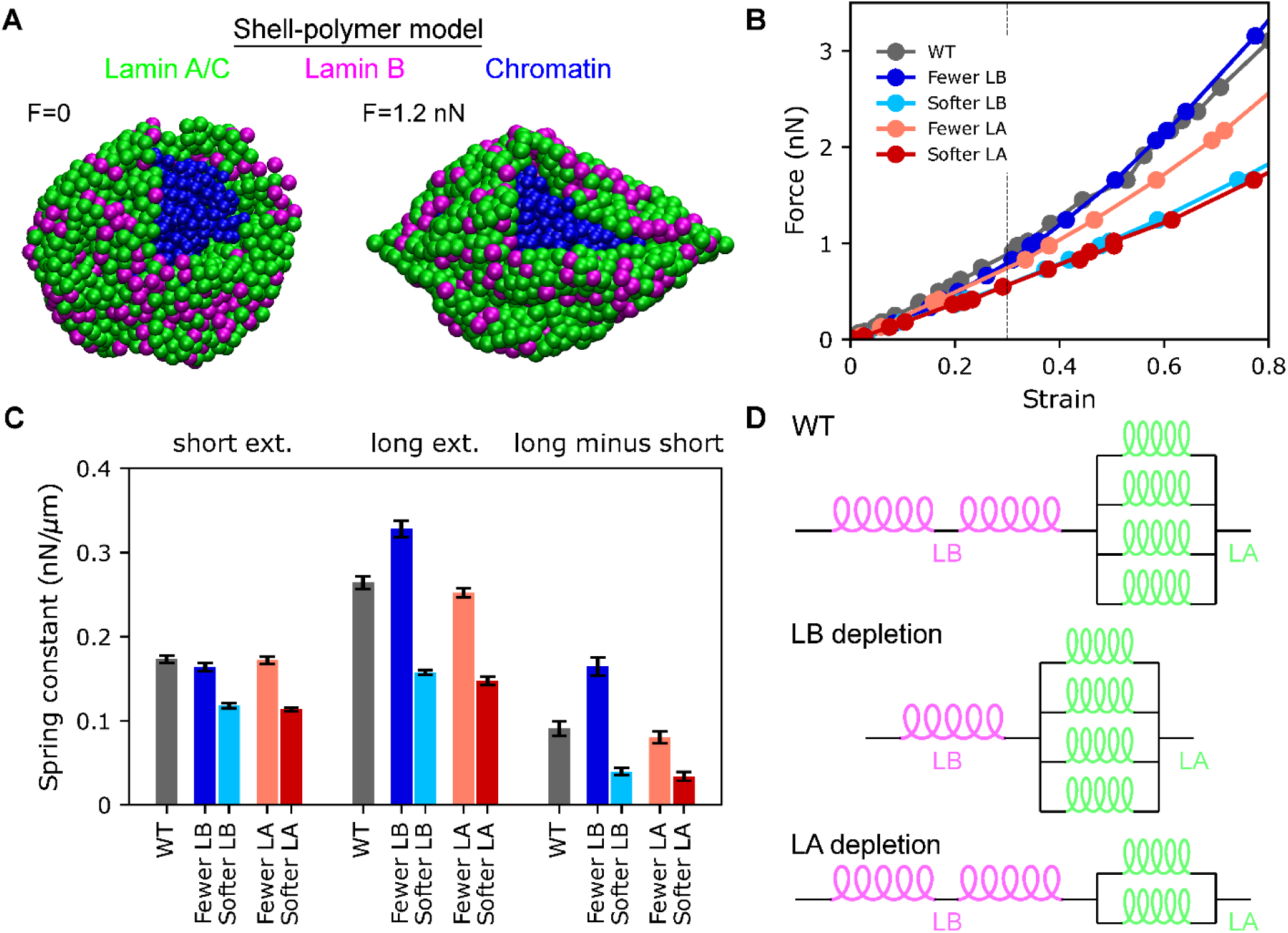
Polymer simulation model of lamina encapsulating chromatin demonstrates differential effects of lamin perturbations. (A) Snapshots of shell-polymer simulations with extensile forces F=0 and 1.2 nN, showing lamin A/C and lamin B subunits in the shell (green and pink, respectively) and a portion of the shell removed to reveal the crosslinked chromatin polymer inside (blue). (B) Force-strain relation for simulations in model with high WT (gray) lamin A (75% lamin A subunits, 25% lamin B subunits). Force-strain relations are also shown for simulations with fewer lamin A or B subunits (light red or dark blue) or softer lamin A or B springs (light blue or dark red). Dotted line indicates 30% strain. Data points represent the average of n≥11 simulations. (C) Bar plot indicating fitted spring constants in the short-extension (left, strain<0.3) and long-extension regimes (middle, 0.3<strain<0.8) and the difference between the two (right). (D) Conceptual model of a lamina composed of lamins A and B. In the model, depletion of lamin B removes soft springs in series (loose pink coils), which stiffens the entire structure, while depletion of lamin A removes stiff springs in parallel (tight green coils), softening the lamina.

For each type of lamina subunit, we considered two perturbations: 1) reduction in the number of subunits, i.e., correlating lamin levels to the number of the nodes in the shell (or area of faces in the network (Shimi *et al*., 2015; Hameed *et al*., 2026)), and 2) softening of the associated springs, i.e., correlating lamin levels to local lamina elasticity (e.g., through the thickness of the fibers formed in the lamina), and thus, node elasticity as in previous simulations (Banigan *et al*., 2017; Stephens *et al*., 2017).

These perturbations to the lamina have unique effects on mechanical response in the simulations in different regimes of extension (**Figure 4B-C and Supplemental Figure 3**). In the base model, with 75% lamin A and 25% lamin B, reducing the number of lamin A or B subunits by 40% results in minimal changes to the short-extension spring constant compared to the wild type scenario (**Figure 4C**). In contrast, reducing the stiffness of the springs associated with either type of lamin subunit by 70% generally resulted in a decrease in short-extension stiffness in the model, depending on the lamin A:B ratio (**Figure 4C and Supplemental Figure 3**). We conclude that changing the material properties of the lamina by reducing the stiffness of nodes within the lamin meshwork can reduce short-extension elasticity, opposite to the effect of changing meshwork architecture.

Next, we observed in the long-extension regime in the model upon changes to lamin A and B nodes and spring constants. Across all three models with different WT levels of lamin B, reducing the number of lamin B nodes by 40% results in ∼20% stiffening of the long-extension spring constant. Reducing the number of lamin A nodes has little effect on long-extension spring constant. However, reducing the spring constants of lamins A and B by 70% decreases the long-extension spring constant in all scenarios (**Figure 4C and Supplemental Figure 3**). The strength of the decrease depends on the composition of the lamina, with nuclei with higher lamin A (or B) being more sensitive to the lamin A (or B) spring constant. Thus, the decreasing the number of lamin nodes and decreasing the stiffness of the springs connecting the nodes have opposing effects on long-extension mechanical response.

Together, these findings illustrate that reductions in lamins that have differential effects on lamina architecture and material properties could result in qualitatively different effects on nuclear mechanical response. The model suggests that lamin B knockdown alters lamina architecture, for instance by reducing the number of nodes, while lamin A knockdown alters the material properties of the network, possibly by altering the stiffness of lamin fibers at the nuclear periphery. Analogously, within the lamina, lamin B may play the role of soft springs in series, while lamin A constitutes a mechanical unit of multiple stiff springs in parallel (**Figure 4D**), leading to the mechanical effects observed in simulations and experiments.

### Nuclear shape and rupture are determined differentially by chromatin and lamin nuclear mechanics regimes

A major function of nuclear stiffness is the maintenance of nuclear shape and integrity, loss of which causes cellular dysfunction (Kalukula *et al*., 2022). To determine how changes in the chromatin-based vs. lamin A/C-based nuclear mechanics regimes affect nuclear shape and integrity, we measured nuclear blebbing through population images and time lapse imaging. First, nuclear shape across the population was measured by the percentage of cells presenting a nuclear bleb and circularity which measures overall nuclear shape. Nuclear blebs were determined as > 1 µm protrusion with decreased DNA density (Bunner *et al*., 2024; Chu *et al*., 2025), and bleb frequency is reported as a percentage (**Figure 5A**). Non-blebbed nuclei were measured for circularity where a perfect circle is 1 and deviation decreases the value (**Figure 5B**).

**Figure 5.**
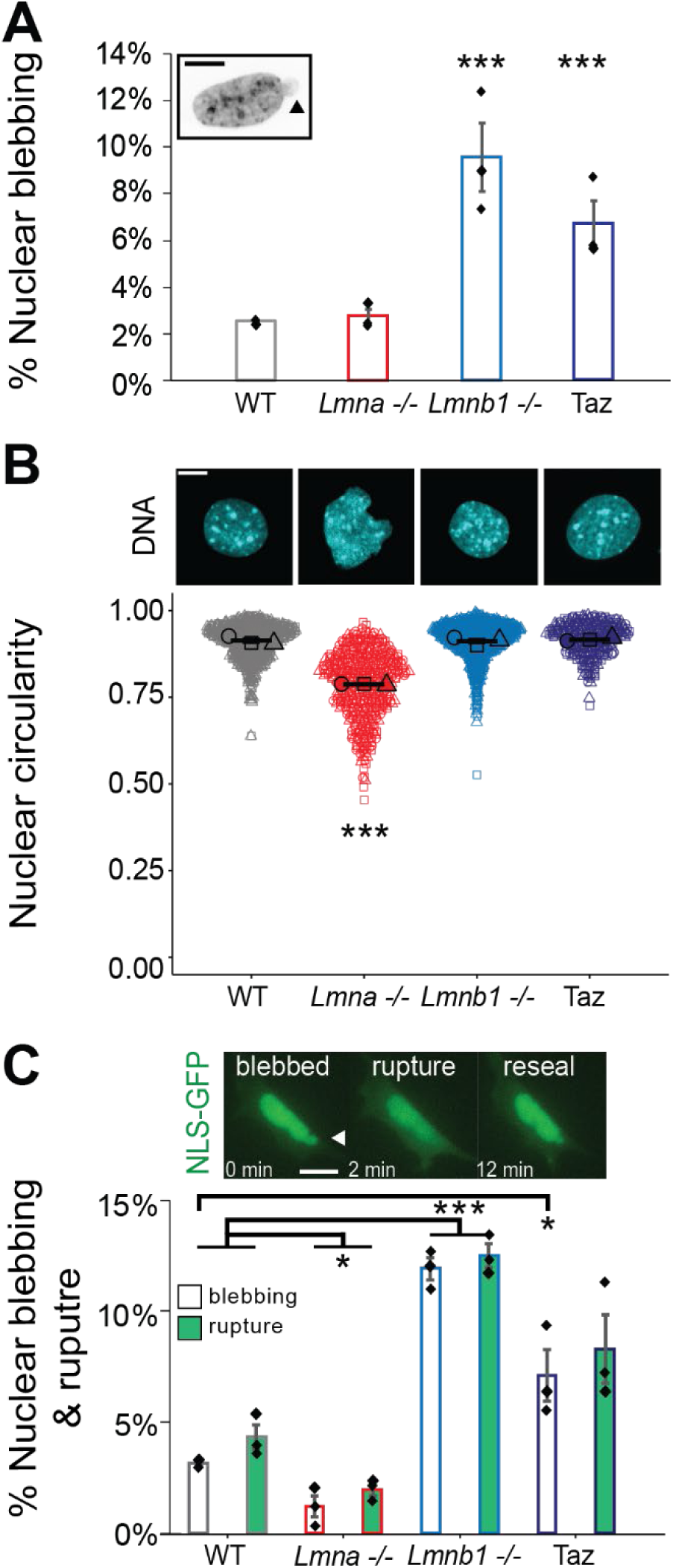
Loss of nuclear stiffness from lamins or chromatin has differential effects on nuclear shape and rupture. Example images and graphs of (**A**) nuclear blebbing percentage and (**B**) circularity of non-blebbed nuclei in static imaging of wild type (WT, gray), lamin A/C knockout (*Lmna-/-*, red), lamin B1 knockout (*Lmnb1-/-*, blue), and loss of facultative heterochromatin (Taz, purple). (C) Example live cell NLS-GFP images depicting a blebbed nucleus, rupture, and post-rupture reseal. Bar graph of the percentage of blebbing (hollow bars) and rupture (solid green bars) during live cell 3-hour imaging for wild type (WT, gray), lamin A/C knockout (*Lmna-/-*, red), lamin B1 knockout (*Lmnb1-/-*, blue), and loss of facultative heterochromatin (Taz, purple). Nuclear blebs denoted by arrow. Biological triplicates each with n > 200 for blebbing and n > 90 nuclei for circularity. For all panels mean and standard error, p values reported as *P<0.05; **P<0.01; ***P<0.001, or ns denotes no significance. Statistical tests are one-way ANOVA with a post-hoc Tukey test. Scale bar = 10 µm.

Loss of lamins A and C in *Lmna-/-* cells resulted in no significant change in the percentage of nuclear blebbing but did result in a significant decrease in nuclear circularity (**Figure 5, A and B**). In agreement with many previous publications (Vargas *et al*., 2012; Hatch and Hetzer, 2016; Stephens *et al*., 2019b), loss of lamin B1 in *Lmnb1-/-* resulted in increased levels of nuclear blebbing, but non-blebbed nuclei maintained circularity. Cells treated with Tazemetostat to decrease facultative heterochromatin also showed increased levels of nuclear blebbing without changing circularity in non-blebbed nuclei, similar to the effects of lamin B1 knockout (**Figure 5, A and B**). Loss of lamin B2 (*Lmnb2-/-*) showed no change in nuclear blebbing (**Supplemental Figure 2**). Thus, loss of all lamin-based strain stiffening in *Lmna-/-* affects nuclear circularity whereas loss of chromatin-based nuclear stiffness in *Lmnb1-/-* and Tazemetostat affect nuclear blebbing.

To measure nuclear blebbing and rupture, we performed time lapse imaging of nuclear localization signal green fluorescent protein (NLS-GFP) in cells (Pho *et al*., 2023). *Lmna-/-* nuclei are abnormally shaped (less circular) and constantly change shape during time lapse movies. Nuclear blebbing decreased significantly from 3.2 ± 0.1 % in wild type to 1.2 ± 0.1 % in *Lmna-/-* nuclei (**Figure 5C**). In agreement with decreased nuclear blebbing, *Lmna-/-* nuclei rarely rupture with an overall significant decrease to 2 ± 0.5 % compared to 4.3 ± 0.9 % nuclear ruptures in wild type (**Figure 5C**). *Lmnb1-/-* nuclei showed a drastic increase in nuclear blebbing and rupture, as blebs account for the large majority of nuclear ruptures. Tazemetostat-induced loss of facultative heterochromatin also increased nuclear blebbing and rupture. This data supports that nuclear ruptures are primarily caused by nuclear blebs, and not overall shape change. Time lapse data reveal that loss lamin-based strain stiffening in *Lmna-/-* decreases nuclear blebbing whereas loss of chromatin-based nuclear stiffness in *Lmnb1-/-* and Tazemetostat increases nuclear blebbing and rupture.

## Discussion

Although lamins have long been associated with nuclear mechanics, lamin B1 has exhibited confounding effects. Initial reports stated that lamin B1 has no mechanical role in the nucleus but does affect morphology (Lammerding *et al*., 2006). Other studies observed that reducing lamin B1 levels could in fact *increase* nuclear stiffness (Shin *et al*., 2013; Swift *et al*., 2013). We have provided novel data detailing that lamin B1 is a critical for heterochromatin maintenance, which in turn determines short-extension nuclear stiffness, shape through nuclear blebbing, and integrity as determined by nuclear rupture. Using lamin knockout cell lines, we confirm that lamin A/C’s mechanical role in the nucleus is to generate stiffness in response to large deformations, whereas lamin B1 primarily weakens the lamina in this regime. In the broader context of previous studies, our data and simulations reveal that lamin B1 may constitute a soft meshwork and depletion of lamin B1 may weaken short-extension nuclear mechanical response through heterochromatin while resisting large deformations by altering the architecture of the nuclear lamina. We reveal the underlying mechanisms for lamin B1’s role in nuclear mechanics, shape, and integrity.

### The mechanical role of lamin B1

Lamin B1’s role in nuclear mechanics and integrity is based on its maintenance of facultative heterochromatin. Our data agrees with a previous publications showing that lamin B1 is essential for facultative heterochromatin maintenance through H3K27me3 (Camps *et al*., 2014; Stephens *et al*., 2018; Vahabikashi *et al*., 2022; Pho *et al*., 2023). The measurements made by micromanipulation demonstrate that loss of lamin B1 is associated with a loss of chromatin-based nuclear mechanics at short extensions, rather than a change in stiffness of the lamina that occurs at longer extensions. This loss of chromatin-based nuclear mechanics can be recapitulated by loss of facultative heterochromatin H3K27me3 through EZH2 inhibitor Tazemetostat (**Figure 1C and 2C**). We previously showed that loss of H3K27me3 via methyltransferase inhibitor DZNep decreases chromatin-based force response without affecting lamin strain-stiffening (Stephens *et al*., 2018; Manning *et al*., 2025). This suggests that EZH2 and H3K27me3 are reliant on lamin B1. This idea is supported by chromatin immunoprecipitation data that H3K27me3 typically enriches lamin associated domains (LADs) associated with lamin B1, and H3K27me3 enrichment at LADs is lost upon loss of lamin B1 in senescence and in lung cancer (Jia *et al*., 2019).

The presence of lamin B1 generally weakens the long-extension lamin strain-stiffening regime. Our data agrees with the previous work establishing that lamin-based nuclear strength is determined by the ratio of lamin A:B (Swift *et al*., 2013; Harada *et al*., 2014). Consequently, lamin B1 depletion can stiffen nuclei, especially in nuclei with small ratios of lamin A:B (Shin *et al*., 2013; Stephens *et al*., 2017). This effect is milder in high lamin A nuclei as shown in Hela nuclei (Stephens *et al*., 2017) and reported in this manuscript (**Figure 2**). Overall, our results are consistent with previous findings that lamin-based nuclear stiffness is well described by the ratio of lamin A:B, with the caveat that overall lamin levels modulate how drastically mechanics change (Swift *et al*., 2013).

Our polymer simulation model suggests an explanation for the confounding finding that removal of a lamin protein (lamin B1) can strengthen the nuclear lamin meshwork. While this physical phenomenon is analogous to removing springs set in series, this simple mode of lamin organization does not capture the mechanics of lamin A/C proteins, which act more like springs in parallel (**Figure 4D**). In our simulation model, removal of lamin B subunits – soft nodes in the network – results in stiffening of the nucleus. In contrast, further softening of the lamin B nodes results in a softer nuclear response. This suggests that, to the extent that lamin B may contribute to mechanics of the lamina, nuclear stiffening upon lamin B1 depletion results from organizational rather than material changes to the lamin meshwork. Consistent with this idea, experiments have observed that lamin B1 knockout changes the structure of the lamin network (Shimi *et al*., 2015; Alabi *et al*., 2025). For example, the mean area per meshwork face increases, which is consistent with the notion of fewer nodes in the network. In contrast, lamin A/C depletion is better recapitulated in the model by a decrease in the spring constant of lamin A subunits. We therefore hypothesize that nuclear softening with lamin A/C depletion is more likely to arise from changes in the elasticity of the structures (fibers) within the lamin A/C network. Intriguingly, the model also suggests that lamin B1 structures should be softer than lamin A/C structures in order to obtain these effects; across all simulated lamina compositions, reducing the number of stiff subunits in the polymer shell had minimal effect on nuclear stiffness. This is consistent with previous findings suggesting that lamin B1 constitutes a deformable network (Nmezi *et al*., 2019), while lamin A/C provides the dominant contribution to the elastic response of the lamina (Swift *et al*., 2013).

### Facultative and constitutive heterochromatin have differing mechanical roles but similar effects on nuclear integrity

The two different types of heterochromatin contribute to nuclear stiffness with one key difference. Loss of facultative heterochromatin, defined by variable silencing of genes dependent on cell type (Lee *et al*., 2020) and tracked via H3K27me3, results in a decrease in the short-extension chromatin-based nuclear spring constant but has no effect on long extension lamin-based strain stiffening. This is a different mechanical outcome than loss of constitutive heterochromatin, defined as heterochromatin consistent across all cell types (Lee *et al*., 2020). Our recent studies report that loss of constitutive heterochromatin marker H3K9me3 decreases both the chromatin-based and lamin-based nuclear stiffness (Manning *et al*., 2025; Bunner *et al*., 2026). Modeling of nuclear mechanics suggests that physical linkages between heterochromatin and the nuclear periphery are crucial for both short-extension (chromatin-based) and long-extension strain stiffening (lamin-based) nuclear mechanical response (Strom *et al*., 2021; Attar *et al*., 2025). The difference is likely because facultative H3K27me3 is not associated with the nuclear periphery to the degree of H3K9me3, which has a prominent enrichment at the periphery, binding to the lamins (Shevelyov and Ulianov, 2019)

While they play slightly differing mechanical roles, loss of either heterochromatin type results in increased nuclear blebbing and rupture, which is known to cause cellular dysfunction (Manning *et al*., 2025). This is interesting because their differing roles suggest that loss of both could additively exacerbate defects in mechanics, morphology, and integrity. Consistent with this notion, loss of both H3K27me3 and H3K9me3 are reported in advanced aging diseases, such as Hutchinson Gilford Progeria Syndrome (HGPS) (Shumaker *et al*., 2006; Stephens *et al*., 2018). Furthermore, while many lamin A/C perturbations induce abnormal nuclear shape but not blebbing, this is not the case with HGPS, which shows drastic nuclear blebbing and ruptures (Kim *et al*., 2021). This points to the mechanistic importance of changes in chromatin modifications in HGPS and other conditions exhibiting nuclear blebbing.

### Nuclear stiffness determines nuclear shape and integrity differentially

Loss of lamin A/C causes loss of strain stiffening but does not result in nuclear blebs. Instead, loss of the lamin regime results in overall loss of nuclear shape. This outcome differs from that of loss of chromatin-based nuclear stiffness, which does result in nuclear blebbing. Consistent with the effects of lamin B1 on chromatin-based nuclear stiffness, lamin B1 loss also induces nuclear blebbing, likely due to the consequent loss of facultative heterochromatin. Below we discuss a possible hypothesis underlying the mechanism by which perturbations to chromatin-based nuclear response to small deformations results in a different nuclear shape outcome than loss of lamin A/C-based nuclear strain stiffening at longer extensions.

We propose that chromatin escapes gaps in the lamina to form a nuclear bleb is a plausible mechanism with previous support (Nmezi *et al*., 2019; Alabi *et al*., 2025). In this model the lamin A/C acts as a resistive meshwork at the periphery containing the chromatin gel. When force is applied to the nucleus, the lamin meshwork equally distributes the stress of chromatin pressing against it to maintain ellipsoidal nuclear shape (Lele *et al*., 2018; Tocco *et al*., 2018). However, upon decompaction, chromatin can escape through a gap in the lamin meshwork, herniate out of the nuclear periphery, and form a nuclear bleb at a defined site. More specifically, chromatin may herniate at a lamin A/C gap that is weaker through chromatin motion pushing driven by active transcription (Berg *et al*., 2023; Prince *et al*., 2025). In this scenario, both stiffness of the nucleus to resist deformation and gaps in the lamin network would dictate nuclear blebbing.

This hypothesis relies on the existence of a strong lamin meshwork to resist outward pressure from the interior chromatin gel, which is compressed by the confining forces of the cytoskeleton (Hatch and Hetzer, 2016; Mistriotis *et al*., 2019; Pho *et al*., 2023; Bunner *et al*., 2026). Complete absence of a resistive lamin meshwork, as in the case of lamin A/C depletion, would lead to a lack of focused herniation through a single large pore. This could result in transient fluctuations and/or larger deformations rather than blebs. In fact, our experimental data confirm this outcome as lamin A/C knockout nuclei display large deformations that are fluid (**Figure 5**). Our data agrees with others that loss of all lamins suppresses nuclear blebbing (Chen *et al*., 2018). Thus, loss of lamin A/C-based nuclear stiffness impacts nuclear shape but suppresses nuclear bleb formation.

In this picture, lamin B1 knockout might alter the configuration of the peripheral meshwork, as suggested by our simulations (**Figure 4**). For instance, the increase in area per face (gap) in the lamina upon lamin B1 knockout (Shimi *et al*., 2015; Alabi *et al*., 2025) could facilitate the emergence of a single dominant herniation, such as a bleb, when the nucleus is mechanically stressed.

Several experimental results support our conceptual model of blebbing by chromatin gapping the lamin A/C meshwork. As noted, increased lamin meshwork size or pore size has been reported for lamin mutants, including lamin B1 (Shimi *et al*., 2015; Alabi *et al*., 2025). Loss of LAP2β has also been shown to increase lamin meshwork pore sizes, resulting in nuclear blebbing (Alabi *et al*., 2025). To further investigate our hypothesis, we propose that the key experiment would be to measure lamin A/C meshwork size in nuclei with chromatin perturbations that cause nuclear blebbing but do not influence lamin levels or lamin mechanics, such as increased euchromatin via valproic acid or decreased heterochromatin by Tazemetostat. This experiment would provide crucial missing evidence for the otherwise enticing hypothesis that flow of chromatin herniating the lamin meshwork serves as a mechanism for nuclear blebbing.

The differences between lamin A/C and B1 continue into nuclear integrity. Loss of lamin A/C abnormal overall nuclear shape results in less nuclear ruptures. Oppositely, loss of lamin B1 due to loss of facultative heterochromatin, and possibly increased lamin A/C gaps, results in nuclear ruptures. We propose the nuclear blebs are essential to nuclear ruptures, while large deformations, such as those in lamin A/C knockout nuclei, are insufficient to increase or induce nuclear ruptures. Previously we showed that > 95% of nuclear blebs result in nuclear rupture (Stephens *et al*., 2019b). This is fundamentally important because nuclear rupture has now been reproducibly shown to cause dysfunction including DNA damage, inflammation, transcription, and cell cycle control (De Vos *et al*., 2011; Helfand *et al*., 2012; Denais *et al*., 2016; Raab *et al*., 2016; Pfeifer *et al*., 2018; Cho *et al*., 2019; Stephens *et al*., 2019b; Pho *et al*., 2023). We recently showed that some of these dysfunctions can create a feedback loop to generate more nuclear blebbing and leading to more dysfunction (Eskndir *et al*., 2025). Thus, the mechanical differences between lamin A/C and B1 fundamentally influence cellular behavior to be healthy or dysfunctional.

## Materials and Methods

### Cell culture

Mouse embryonic fibroblasts (MEFs) were previously described (Shimi *et al*., 2008; Kim *et al*., 2011; Stephens *et al*., 2018; Vahabikashi *et al*., 2022). MEF wild-type (WT), lamin A/C knockout (*Lmna -/-*), lamin B1 knockout (*Lmnb1 -/-*), and lamin A knockdown (LA KD) were cultured in a 60 mm culture dish in DMEM (Corning, 45000-304) supplemented with 10% fetal bovine serum (FBS; Cytiva HyClone, SH3007103) and 1% Penicillin/streptomycin (PS; Corning, MT30002CI), incubated at 37°C and 5% CO2. After reaching 80-90% confluency, cells were passaged by being trypsinize with 0.25% Trypsin, 0.1% EDTA without sodium bicarbonate (Corning, MT25053CI), replated, and diluted with DMEM. We thank Zhang lab and Goldman lab for lending us the *Lmna-/-* and *Lmnb1-/-* cell lines (Shimi *et al*., 2015).

### Drug treatments

Cells were treated with EZH2 inhibitor tazemetostat (Taz; MedChemExpress, HY-13803) at 10-20µM for 16 hours.

### Live-cell time-lapse fluorescence imaging and analysis

As previously established, we used MEF NLS-GFP stable cell lines to quanitfy nuclear shape and rupture (Pho *et al*., 2023). Imaging was performed using a Nikon Instruments Ti2-E microscope, Orca Fusion Gen III camera, Lumencor Aura III light engine, TMC CleanBench air table, with objective lens Plan Abo Lambda 40x air (N.A. 0.75, W.D. 0.66, NMRH00401). Images were acquired using the Nikon Perfect Focus System and with Nikon Elements software. Cells were kept alive throughout time-lapse imaging using Okolab heat at 37 °C, supplemental humidity, and a 5% CO2 stage top incubator (H301). Images were captured at 30ms exposure time, 12-bit depth, and 3% blue fluorescent light (475 nm) power. Time lapse images were analyzed by visually tracking bleb formation and rupture throughout the movie. To analyze, the total number of blebbed nuclei, ruptured nuclei, and number of nuclei per field of view were recorded. In order to calculate blebbing and rupture percentages, the number of blebbed or ruptured nuclei was divided by the number of total nuclei per field of view. Then, this average was used to compare differences between treatment conditions and cell perturbations.

### Immunofluorescence (IF)

Cells were fixed with 4% paraformaldehyde (PFA; Electron Microscopy Sciences, 50-980-487) in phosphate buffered saline (PBS; Corning, 45000-446) for 15 minutes then washed 3 times in PBS for 5 min. Cells were permeabilized with 0.1% Triton X-100 (VWR Life Science, 9002-93-01) in PBS for 15 min and washed with 0.06% Tween-20 (Promega, H5152) in PBS for 5 min. Cells were then washed again 3 times in PBS for 5 min. Cells were then blocked for 1 h at room temperature with 2% bovine serum albumin (BSA; Fisher BioReagents, BP1605-100) in PBS followed by overnight staining at 4 °C with primary antibodies. Primary antibodies were diluted in the blocking solution at the following concentrations: H3K27me3 (1:1000 Cell Signaling Technology, 9733s), lamin A/C (1:250 Cell Signaling Technology, 4777s), lamin B1 (1:1000 abcam, ab16048), H3K9me3 (1:1000 abcam ab8898), LBR (1:500 abcam, ab232731), LAP2β (1:500 Thermo Fisher, PA5-52519), lamin B1 (1:1000 abcam, EPR9701[B]). Cells were washed 3 times with PBS. Cells were incubated on a nutating mixer in the dark with the following secondary antibodies at a 1:1000 dilution in BSA blocking solution: Alexa Fluor 555 anti-mouse IgG (1:1000 Cell Signaling Technology, 4409s), Alexa Fluor 647 anti-mouse IgG (1:1000 Cell Signaling Technology, 4410s), Alexa Fluor 555 anti-rabbit IgG (1:1000 Cell Signaling Technology, 4413s), Alexa Fluor 647 anti-rabbit IgG (1:1000 Cell Signaling Technology, 4414s). Cells were then washed with PBS 3 times for 5 min. Cells were stained with Hoechst 33342 (1:10,000, Invitrogen H3570) in PBS for 15 min. Cells were washed a final time with PBS 3 times for 5 min.

### Fluorescence imaging and analysis

Cells were imaged using a Nikon Instruments Ti2-E microscope with Crest V3 Spinning Disk Confocal, Hamamatsu Orca Fusion Gen III camera, Lumencor Aura III light engine, TMC CleanBench air table, with 40x air objective (N.A. 0.95, W.D. 0.66, NMRH00401). Images were acquired with a 12-bit camera using the Nikon Perfect Focus System and with Nikon Elements software. Images were taken at 0.5 µm z-steps over 4.5 µm. For intensity measurements, greater than 100 nuclei were assayed and regions of interest were selected for analysis in NIS-Elements. Background signal was subtracted from each image before measurements were taken. Relative intensity values were exported to Automated Results and Measurements, then to an Excel sheet for analysis. The average mean intensity of a given fluorophore was calculated between conditions. To measure circularity, ROIs were copied to Binary, then circularity values were exported to Automated Results and Measurements and finally exported to an Excel sheet for analysis.

### Fluorescence imaging and analysis of nuclear periphery

Cells were imaged using a Nikon Instruments Ti2-E microscope with Crest V3 Spinning Disk Confocal, Hamamatsu Orca Fusion Gen III camera, Lumencor Aura III light engine, TMC CleanBench air table, with 100x oil immersion objective (N.A. 1.45, W.D. 0.13 mm, MRD71970). Images were acquired with a 12-bit camera using the Nikon Perfect Focus System and with Nikon Elements software. Images were captured using 13 Z-stacks at a 0.5 µm step. The brightest plane was used for analysis. Using NIS-Elements, the average background fluorescence was subtracted via an ROI with no cells. Individual nuclei were then auto selected as ROIs in binary. On the same nucleus, the ROI was copied and eroded by 8 pixels (0.5 µm) for LAP2β and LBR and 15 pixels (1 µm) for H3K9me3. The measurements of the two ROIs were exported to Excel. The average intensity and ROI size were subtracted from the two ROIs to provide peripheral intensity measurements. For LAP2β and LBR, the average intensity and ROI size of the periphery were multiplied to measure the total fluorescence of the periphery. For H3K9me3, the H3K9me3 fluorescence at the periphery was measured relative to the total fluorescence of H3K9me3 in the nucleus.

### Micromanipulation force measurement of an isolated nucleus

As previously described (Currey *et al*., 2022; Manning *et al*., 2025), MEF cells were grown in a micromanipulation well and treated as described above. Before isolation, nuclei were treated with actin depolymerization drug latrunculin A (Cayman Chemical;10 µM) concentration for 30 minutes to aid nuclear isolation. Nuclei were isolated from living cells using a spray micropipette containing Triton X-100 (0.05%) in PBS. A pull micropipette was used to grab the nucleus, while the opposite end of the nucleus was grabbed by a precalibrated force micropipette and suspended in preparation for force-extension measurements. The pull pipette was moved 100 nm/s to extend the nucleus while both the pull and force pipettes’ positions were tracked at 15 frames per second. This data was exported to Excel for analysis. Nucleus extension was measured by tracking the change in distance (µm) between the pull and force micropipettes. Force (nN) was measured by using Hooke’s law F = kx, where x is the deflection distance (µm) of the force pipettes from its initial position multiplied by k, the precalibrated bending modulus (nN/µm). This force pipette was precalibrated to a set range of 1.2–2 nN/µm bending modulus. In Excel, the force vs. extension is plotted. The slope of the force vs. extension plot provides the spring constant (nN/µm) for the short chromatin-dominated regime (<3 µm) and long-extension lamin A-dominated strain-stiffening regime (>3 µm). Each nucleus is force-extension measured three times and averaged to provide one composite data point. The long-regime spring constant minus the short-regime spring constant provides the measure of lamin-based strain stiffening.

## Statistics

Each figure legend specifies which statistical test was used on the data. A one-way ANOVA with post-hoc Tukey test was run on data with multiple conditions for Figures 1,2,3,5 and Supplemental Figure 1. Data sets that consist of only two conditions or could not be reliably tested for normality due to limited numbers of replicates (data sets with only 3 replicates per condition) were assessed for statistical significance via a Student’s t-test. Two-tailed unpaired Student’s t-tests were run on Supplemental Figure 2.

### Simulations

Brownian dynamics simulations were performed by adapting a previously developed polymer-shell model (Banigan *et al*., 2017; Stephens *et al*., 2017; Strom *et al*., 2021). The simulation code is freely and publicly available on GitHub (https://github.com/ebanigan/shell-polymer). “Wildtype” simulated nuclei consisted of 1333 lamin subunits of diameter *a_sh_ =* 0.71 μm composing a polymer shell of radius *R* = 10 μm representing the lamina and 552 chromatin subunits of diameter *a_poly_ =* 0.71 μm composing the crosslinked polymer chain representing chromatin. At the start of each simulation, a fixed percentage of lamin subunits were designated as lamin B (75% in WT in the main text and **Figure 4**, with lower levels shown in **Figure S2**). Simulations with reduced levels of lamin A or lamin B contained 40% fewer of the perturbed subunit in the shell, consequently reducing the total number of lamina subunits. Each shell subunit was connected by springs to 4 ≤ *z* ≤ 8 nearest neighbor shell subunits, with a mean connectivity of <*z*> ≈ 4.5. Springs connecting two lamin A subunits had spring constant *k_A_* = 0.8 nN/μm, while those connecting two lamin B subunits had spring constant *k_B_* = 0.2 nN/μm in WT simulations. Springs connecting a lamin A subunit to a lamin B subunit had spring constant *k_AB_* = 2*k_A_k_B_*/(*k_A_*+*k_B_*). In simulations with reduced stiffness of lamin A or lamin B, *k_A_* or *k_B_* were reduced by 70% and *k_AB_* was altered accordingly. The chromatin polymer was a linear polymer connected by springs, crosslinked by an additional *N_c_* = 55 springs, and linked to subunits in the shell by *N_L_* = 40 springs; these springs had spring constant *k*_chr_ = 1.6 nN/μm. Extensile force, *F*, was exerted across the nucleus along the x-axis by exerting force at each of the two poles of the shell. All subunits exerted short-ranged repulsive, excluded-volume-like interactions each other via harmonic potentials (see (Strom *et al*., 2021)). All subunits were subject to thermal noise and obeyed the overdamped Langevin equation. Equations of motion (see (Banigan *et al*., 2017)) were evolved by an Euler algorithm (Allen and Tildesley, 1989) with timestep *dt* = 0.0005.

## Acknowledgements

We would like to thank the Zhang, Reddy, and Goldman labs for sharing lamin knockout and knockdown cell lines. We thank Aykut Erbas for helpful discussions as well as lab mates Madeleine Clark, Antonela Losada, Katie Lin, and Nickolas Borowski for providing support and feedback on the manuscript.

## Funding

This work was supported by NIH NIGMS grant Maximizing Investigators’ Research Award R35GM154928 obtained by ADS. EJB acknowledges support from NSF MCB 2044895.

## Data availability

All raw data that is available https://doi.org/10.6084/m9.figshare.33106643. Image data set can be made available upon request.

## Competing interests

The authors declare no competing interests.

## Notes

https://doi.org/10.6084/m9.figshare.33106643

### Competing Interest Statement

The authors have declared no competing interest.

