## Supplemental Figures 1-3 for "Lamin B1 affects nuclear shape and integrity through chromatin stiffness and not lamin stiffness"

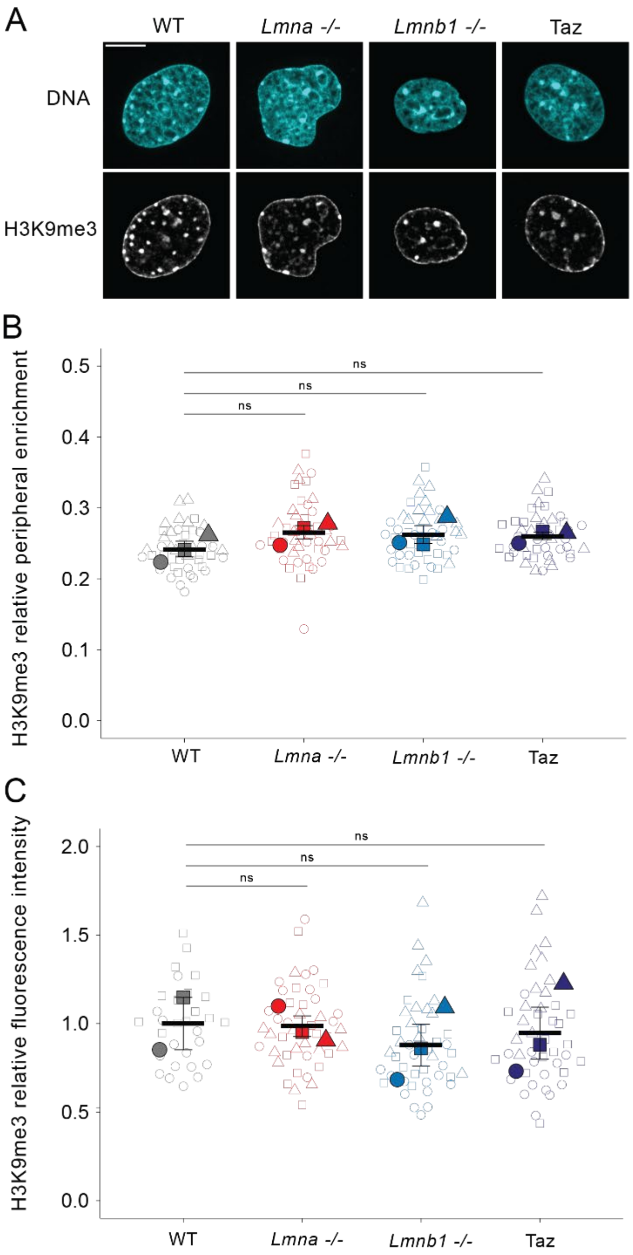


**Supplemental Figure 1. H3K9me3 immunofluorescence periphery enrichment and whole nucleus levels are maintained.** (A) Example confocal images of DNA (Hoechst, cyan) and constitutive heterochromatin marker H3K9me3 in wild type (WT), lamin A/C knockout (*Lmna-/-* , red), lamin B1 knockout (*Lmnb1-/-*, blue), and loss of facultative heterochromatin (Taz, purple). Superplot graphs of constitutive heterochromatin marker H3K9me3 relative (B) peripheral enrichment and (C) overall nuclear intensity in wild type (WT), lamin A/C knockout (*Lmna-/-* , red), lamin B1 knockout (*Lmnb1-/-*, blue), and loss of facultative heterochromatin (Taz, purple). Biological triplicates each with n = 15 nuclei. For all panels mean and standard error, p values reported as *P<0.05; **P<0.01; ***P<0.001, or ns denotes no significance. Statistical significance was determined by one-way ANOVA with a post-hoc Tukey test. Scale bar = 10 μm.

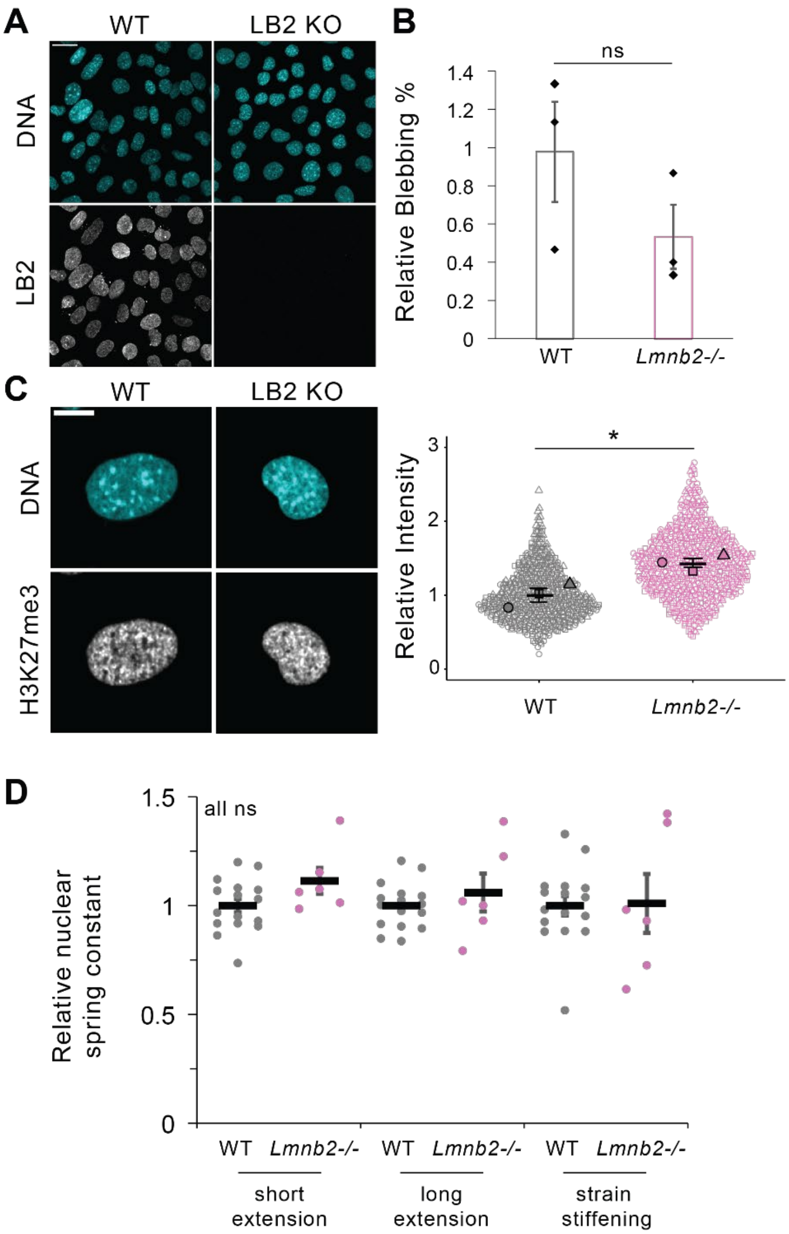


**Supplemental Figure 2. Lamin B2 knockout does not alter nuclear blebbing or micromanipulation nuclear spring constant.** (A) Example images of DNA (Hoechst, cyan) and lamin B2 (gray scale) in wild type and lamin B2 knockout *Lmnb2-/-*. (B) Bar graph of relative nuclear blebbing in wild type (WT, gray) and lamin B2 knockout (*Lmnb2-/-* , pink). Biological triplicates each with n > 300 nuclei each. (C) Example images and superpolt of DNA and facultative heterochromatin marker H3K27me3 in wild type (WT, gray) and lamin B2 knockout (*Lmnb2-/-* , pink). Biological triplicates each with n =200 nuclei each. Dual micropipette micromanipulation force vs. extension relative nuclear spring constants for short extensions (< 3 µm), long extension (> 3 µm), and strain stiffening for wild type (WT, gray, n = 19 nuclei from Figure 2) and lamin B2 knockout (*Lmnb2-/-* , pink, n = 6 nuclei with 26 measurements total). For all panels mean and standard error, p values reported as *P<0.05; **P<0.01; ***P<0.001, or ns denotes no significance. Statistical significance was determined by two-tailed unpaired Students' t-test. Scale bar = 10 µm.

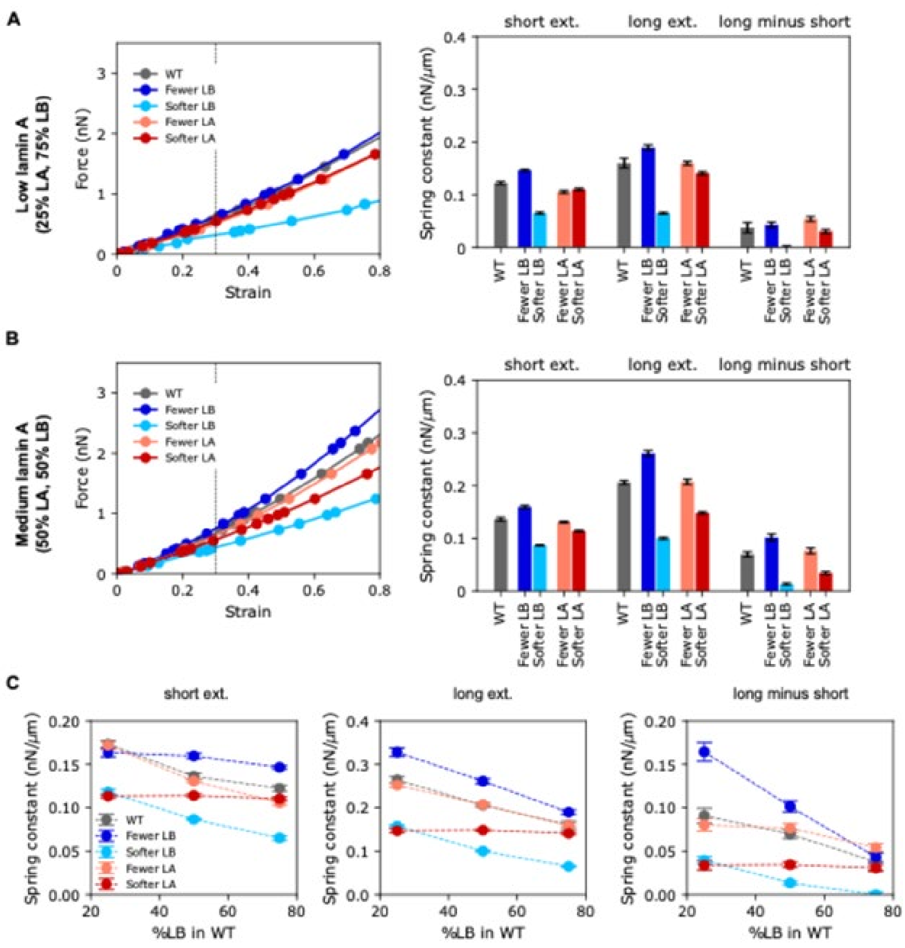


**Supplemental Figure 3. Force response for simulation models with different levels of lamins A and B.** (A-B) Force-strain relations (left) and bar plots of fitted spring constants (right) for simulation models with low (A) or medium (B) levels of lamin A in WT (25% or 50% lamin A, respectively), along with perturbations. (C) Nuclear stiffness in short- and long-extension regimes (left and middle) depending on composition of the lamina in simulations, denoted by percentage of subunits that are lamin B in WT. Rightmost plot depicts the difference in long- and short-extension spring constants.
